# The emergence of a collective sensory response threshold in ant colonies

**DOI:** 10.1101/2021.10.30.466564

**Authors:** Asaf Gal, Daniel J. C. Kronauer

**Affiliations:** Laboratory of Social Evolution and Behavior, The Rockefeller University, 1230 York Avenue, New York, NY 10065

## Abstract

The sensory response threshold is a fundamental biophysical property of biological systems that underlies many physiological and computational functions, and its systematic study has played a pivotal role in uncovering the principles of neural computation. Here, we show that ant colonies, which perform computational tasks at the group level, have emergent collective sensory response thresholds. Colonies respond collectively to step changes in temperature and evacuate the nest during severe perturbations. This response is characterized by a group-size dependent threshold, and the underlying dynamics are dominated by social feedback between the ants. Using a binary network model, we demonstrate that a balance between short-range excitatory and long-range inhibitory interactions can explain the emergence of the collective response threshold and its size dependency. Our findings illustrate how simple social dynamics allow insect colonies to integrate information about the external environment and their internal state to produce adaptive collective responses.

## Introduction

Sensory thresholding is one of the most fundamental and well-studied computational primitives performed by organisms, where a perceived level of sensory input is compared with an internal variable to generate a binary neural, physiological or behavioral response. Organisms rely on sensory thresholds to perform critical functions such as responding to threats, detecting prey and making decisions^1,2^. Sensory thresholds also play important roles as components of more complex tasks such as foraging, navigation, recognition and communication^3–5^. The methodical study of sensory thresholds is a cornerstone of modern neuroscience, and has played a key role in connecting cognitive, behavioral and computational phenomena to neural and biophysical mechanisms^6,7^. At the computational level, thresholds are understood as decisions that optimize costs and benefits associated with responding or not responding in a specific context, and are analyzed using signal detection theory^8^. Mechanistically, thresholds in biological systems almost always emerge from a balance between two opposing forces: exciting and restoring electrical currents in excitable membranes^9^, excitatory and inhibitory neurons in neural networks^10^, or inter-region connections at the level of the entire brain^11^. Similar dynamical phenomena are also abundant in non-neural systems such as the immune system, cellular and microorganismal behavior, or intracellular signaling networks^12–15^.

Sensory response thresholds play an important part in the organization of social insect colonies, which process information and perform cognitive-like functions at the group level^16–19^. The distribution of individual response thresholds in a colony gives rise to behavioral differentiation and division of labor^20–23^, a hallmark of insect societies. Moreover, studies have demonstrated that, in turn, the sensory threshold of colony members can be modulated by their social environment^24,25^, suggesting that threshold dynamics could underlie the adaptation and reconfiguration of collective behavior. However, it is unclear whether sensory thresholding is by itself a computational primitive at the level of the colony, and if so, how it emerges out of the complex interaction network between the individuals in a colony. Colonies of ants and bees collectively perceive and asses their sensory environment and act upon these assessments in a coordinated manner in choice contexts^26–29^. Therefore, it is reasonable to hypothesize that ants can coordinate their behavior in response to a sensory input to create a colony-level thresholded response. Mechanistically, the concept of an ant colony as an excitable system governed by exciting and inhibiting interactions between the ants has been proposed to explain activity waves and temporal oscillations, in direct analogy with neural network dynamics^30–33^. In principle, similar interactions could also give rise to collective behavioral thresholds.

The analogies and parallels between single-animal and group-level computations have inspired a long line of experimental and theoretical work^29,31,34,35^. Yet, our formal understanding of how collective information processing emerges from group dynamics still lags behind our understanding of neural computation. This is mostly due to the lack of convenient experimental paradigms for relating simple, controlled sensory environments to precise measurements of individual and collective responses. Establishing an experimental paradigm for a systematic study of group-level sensory thresholds can therefore contribute to a formal description of emergent collective computation. Such a paradigm should allow to robustly relate features of controlled sensory environments to the behavioral response of the colony and its members. Here, we study the emergence of a sensory response threshold in colonies of the clonal raider ant *Ooceraea biroi*. The clonal raider ant is an attractive model organism for these kinds of experiments^21,36–38^. It provides unparalleled experimental control over the size and composition of colonies, as well as over the genotype and age of each individual ant in the colony, thus allowing standardization of colony features between experimental replicates. We use a custom setup for controlling the thermal environment of the ants and to measure the behavioral responses of colonies to temperature changes, demonstrating that the collective response is indeed characterized by an emergent sensory response threshold. We then combine experimental manipulations of colony size with mathematical modeling of colony dynamics to investigate the relationship between the collective threshold and the underlying interactions between the individual ants.

## Results

### Ants respond to increasing temperature by evacuating the nest

Controlling the sensory environment of a group of freely behaving animals is challenging. We decided to use thermosensation as the sensory modality, because temperature is a scalar property of the environment, and its sensation is minimally dependent on the position and specific behavior of the individual. To study how ants respond to a temperature increase, we developed a behavioral arena in which the ground temperature is controlled by an array of thermo-electric components (Figure S1, Materials and Methods). A thin layer of plaster of Paris constitutes the surface of the arena. This setup allows us to rapidly change the arena’ s ground temperature (Figure S2), while maintaining ground moisture level constant (Figure S3). In all of the following experiments, we placed *O. biroi* colonies of variable sizes in the arena. Unlike most other ants, *O. biroi* is queenless, and all experimental colonies were composed of workers and larvae in a 2:1 ratio. Adult ants were approximately 1 month old, and larvae were 6-7 days old. *O. biroi* reproduces asexually and clonally, providing precise experimental control over an individual’ s genotype. We standardized genotypes by sourcing all individuals from the same clonal line and stock colony (Materials and Methods). At baseline, the set value of the ground temperature controller was 26°C. When ants are placed in the arena, they quickly create a nest by settling around a brood pile (Figure 1A, inset). Typically, a few scouts explore the arena, while most ants remain inside the nest (Figure 1A-B). After a settling period of about 48 hours, we studied how colonies respond to temperature changes by subjecting them to a sequence of perturbation events. In each perturbation, the set temperature was abruptly increased to a higher set value for a period of 15 minutes. Perturbations were spaced by intervals of two hours to allow the ants to resettle. When the perturbation temperature was relatively high, the colonies typically responded with a stereotypical coordinated evacuation of their nesting site. Figure 1B-E and Video 1 show a representative example of a colony of 36 workers and 18 larvae responding to a 40°C perturbation. Following the temperature increase, the ants gradually get excited and increase their activity levels (Figure 1C). After some delay, the colony initiates an ordered evacuation in which all ants leave the nest in a column (Figure 1D-E, Video 1). Because temperature is increased evenly across the entire arena, the ants remain in a high activity ‘ explorative’ state following the initial response. Depending on the perturbation temperature, this state can be highly organized (for relatively low temperatures, Figure 1F), or manifest as a more chaotic and disorganized collective pattern (for high temperatures, Figure 1G). Once the temperature returns to baseline, the ants slowly relax and reform the nest cluster (Figure 1H-I).

**Figure 1:**
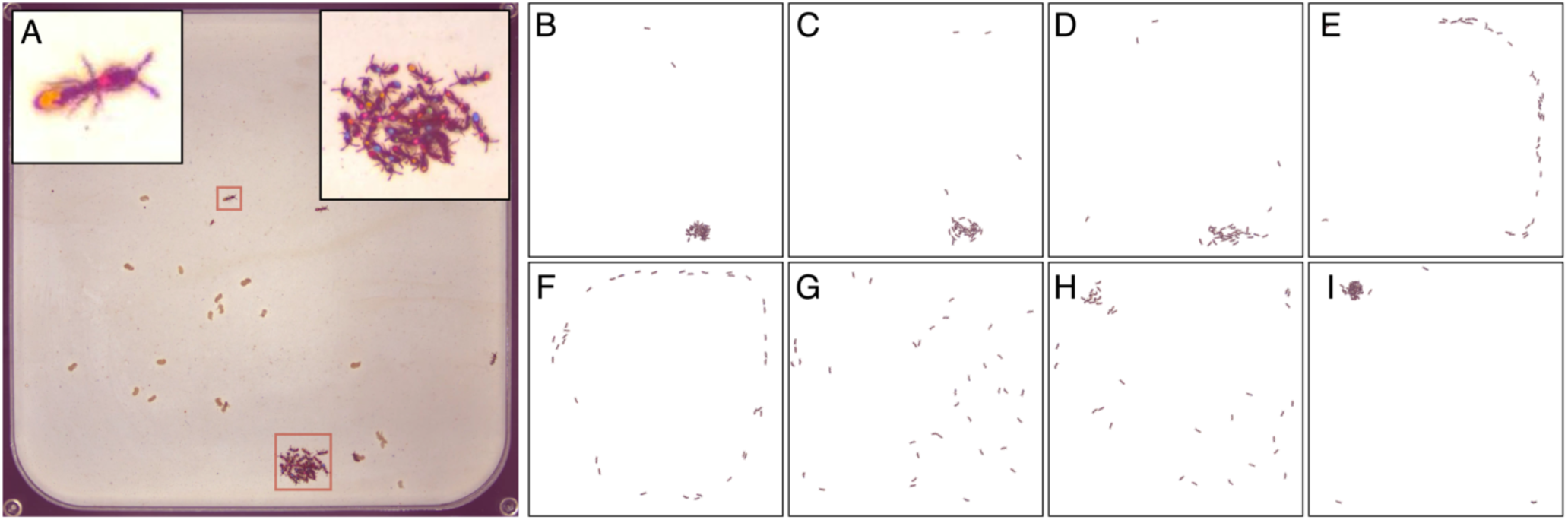
The response of an ant colony to a step temperature perturbation. (**A**) A snapshot from a raw experimental video, showing a colony of 36 ants on a temperature-controlled plaster of Paris arena (Materials and Methods; Figures S1 and S2). Each of the ants is marked with a unique combination of color tags to allow for individual behavioral tracking. The arena is confined by a black metal frame heated to 50°C (Figure S2). The ants form a nest (red square at the bottom and inset at top right), with a few scout ants exploring the arena (top red square and inset at top left). The light brown objects in the arena are food items. (**B-I**) Snapshots depicting the typical dynamics of the response of a colony to a strong temperature perturbation. Images are processed by removing the background for visual clarity. (**B**) *Baseline state*. Before the onset of the perturbation, most ants reside in the nest, with few scout ants exploring the arena. (**C**) *Excitement*. Following the onset of the perturbation, the ants first respond by increasing their activity level around the nest. (**D**) *The onset of emigration*. After a delay that lasts up to a few minutes, the ants suddenly begin to leave the nest in a well-defined direction. (**E**) *Full emigration*. The colony forms a well-organized emigration column. (**F**) *Stable emigration*. (**G**) *Disordered perturbed state*. In some case, especially under high temperature perturbations, the organized emigration column breaks, and the colony enters into a high-activity, swarm-like state, where the movements of the ants are only weakly correlated. (**H**) *Relaxation*. Following the return of the temperature to baseline, the ants slowly relax and begin to reform the nest, possibly in a different location. The relaxation process can take up to one hour to complete. (**I**) *New baseline*. The colony has fully returned to its baseline relaxed state.

### The colony response to temperature perturbations is collective

Escape, or place-change behavior in response to changes in temperature or other environmental parameters is a ubiquitous behavior that is often studied in the context of sensory decision-making^1,39–41^. Solitary animals make such decisions independently, and the correlations between individuals is generally low, both in the decision itself (whether to leave or not) and in the timing and direction of leaving. In contrast, the response of clonal raider ants to the temperature increase seemed highly coordinated, both in time and in space. To quantify the collectivity in this response, we performed an experiment to measure the coordination and correlation between the responses of individual ants. We perturbed 3 colonies of 36 ants and 18 larvae with a sequence of 24 temperature perturbations of 15min duration, each with an amplitude of 33°C, which does not produce a robust evacuation response. Ants in these colonies were marked with unique combinations of color tags, and were individually tracked using a custom software^42^. From the tracking results, the nest location before each perturbation was determined (Materials and Methods). We defined a circle with a 15mm radius around the nest (around twice the typical radius of the nest blob, Figure 2A). The binary response 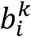 of the *i*-th ant to the *k*-th perturbation was defined as 1 if she exited the nest circle at some point during the time of the perturbation and remained outside for a duration of at least 30 seconds, and as 0 in case she did not. Ants that were outside the nest circle at the beginning of the perturbation were treated as missing values. For each of the 3 colonies, we found that the distribution of average responses across ants for each perturbation is bimodal (Figure 2B), suggesting the decisions to leave the nest are correlated between ants. To test this, we calculated the correlation between the response of each pair of ants in the colony and compared the distribution of these values to a null distribution, constructed by randomly shuffling the individual responses of the ants between perturbation events (Figure 2C, Materials and Methods). These two distributions differ significantly (average pairwise correlation value of 0.396, p < 10^−6^, Student’ s two-sided t-test), showing that the decisions of ants in the colony are indeed highly correlated.

**Figure 2:**
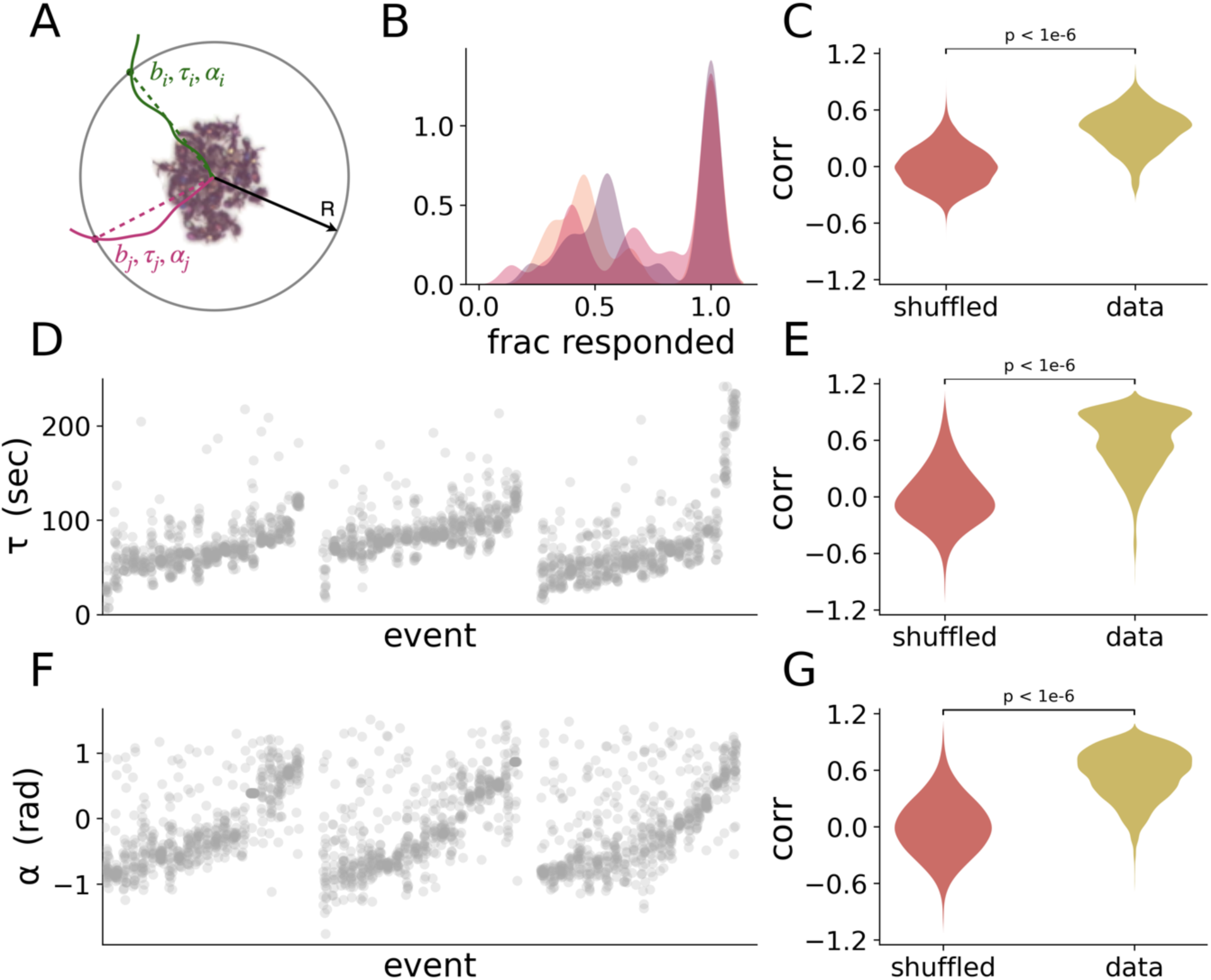
Ants respond collectively to temperature perturbations. (**A**) Measures of individual responses. We define a circle of radius R=15mm around the location of the nest. A schematic drawing of the trajectories of two ants is depicted in green and pink. For each ant, we record the binary response (*b*), the response direction (*α*), and the response latency (*τ*) as her first crossing of that circle for a duration longer than 30 seconds, as explained in the text and the Methods section. (**B**) Distribution of the average binary response across ants. Each distribution is estimated from one colony perturbed with a sequence of 24 perturbations of 33°C. (**C**) Pairwise correlations between the binary responses of ants in **B**, compared to correlations in shuffled responses. Shuffled responses are generated by shuffling the binary responses of each ant to all the perturbations independently of other ants in its colony, therefore eliminating any correlation. The real distribution is composed of 1890 correlation values, produced from the responses of 108 ants from 3 colonies. The null distribution is composed of 189,000 correlation values, produced from 100 independent shuffles of the responses. Statistical significance was estimated using a Student’ s two-sided t-test. (**D**) Scatter plot depicting the distributions of individual response latencies, from 3 colonies perturbed with a sequence of 24 perturbations of 40°C. Each column represents the responses of ants from a single colony to one perturbation. The events are sorted first by colony and then by the average individual response latency in each event. (**E**) Pairwise correlations between the binary responses of ants in **D** compared to a null distribution generated in the same way as in **C**. (**F-G**) Plots as in **D-E**, but for the individual response directions.

To assess whether the ants are also correlated in the timing and direction of their response, we repeated the experiment with 3 additional colonies of the same size and composition, using the same protocol. However, this time we subjected the colonies to stronger perturbations of 40°C, which generally produce a robust collective nest evacuation. We defined the individual response as before, and also measured the response delay 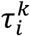 as the time elapsed between the onset of the *k*-th perturbation and the time the *i*-th ant crossed the circle, and the response direction 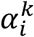 as the angle between the point of crossing and the line connecting the nest center and the center of the arena (Figure 2A). Plotting the distributions of individual response latencies across perturbation events, we found that the variability between individual response latency in the same event is lower than the variability between events (Figure 2D), suggesting that ants are coordinated in the timing of their response. We showed this formally by comparing the distribution of pairwise latency correlations to a null distribution in the same way as before (average pairwise correlation value of 0.585, p < 10^−6^, Student’ s two-sided t-test; Figure 2E). We then repeated the same analysis for the response direction (Figure 2F-G), showing that ants coordinate their response also spatially (average pairwise correlation value of 0.555, p < 10^−6^, Student’ s two-sided t-test).

### The collective response is characterized by a size-dependent threshold

To better understand how the collective response depends on the amplitude of the perturbation, we performed an experiment with 10 colonies of 36 workers and 18 larvae each, and subjected each colony to a sequence of perturbations of variable amplitude, ranging from 28°C to 45°C. We defined the collective state of the colony as the fraction of ants outside of the nest in any given frame (Figure 3A). We defined the binary collective response as 1 if a quorum of at least 90% of the colony was outside the nest for at least 30 consecutive seconds at some point during the time of the perturbation, and 0 otherwise. For each perturbation temperature, we computed the probability of a collective response across all colonies, resulting in a sigmoidal psychometric-like response curve typical for systems with a noisy threshold response (Figure 3B). Using a logistic regression model (Materials and Methods), we estimated the threshold *θ* to be 34.12°C for the colonies in this experiment, with a 95% confidence interval of 33.3°C to 34.8°C.

**Figure 3:**
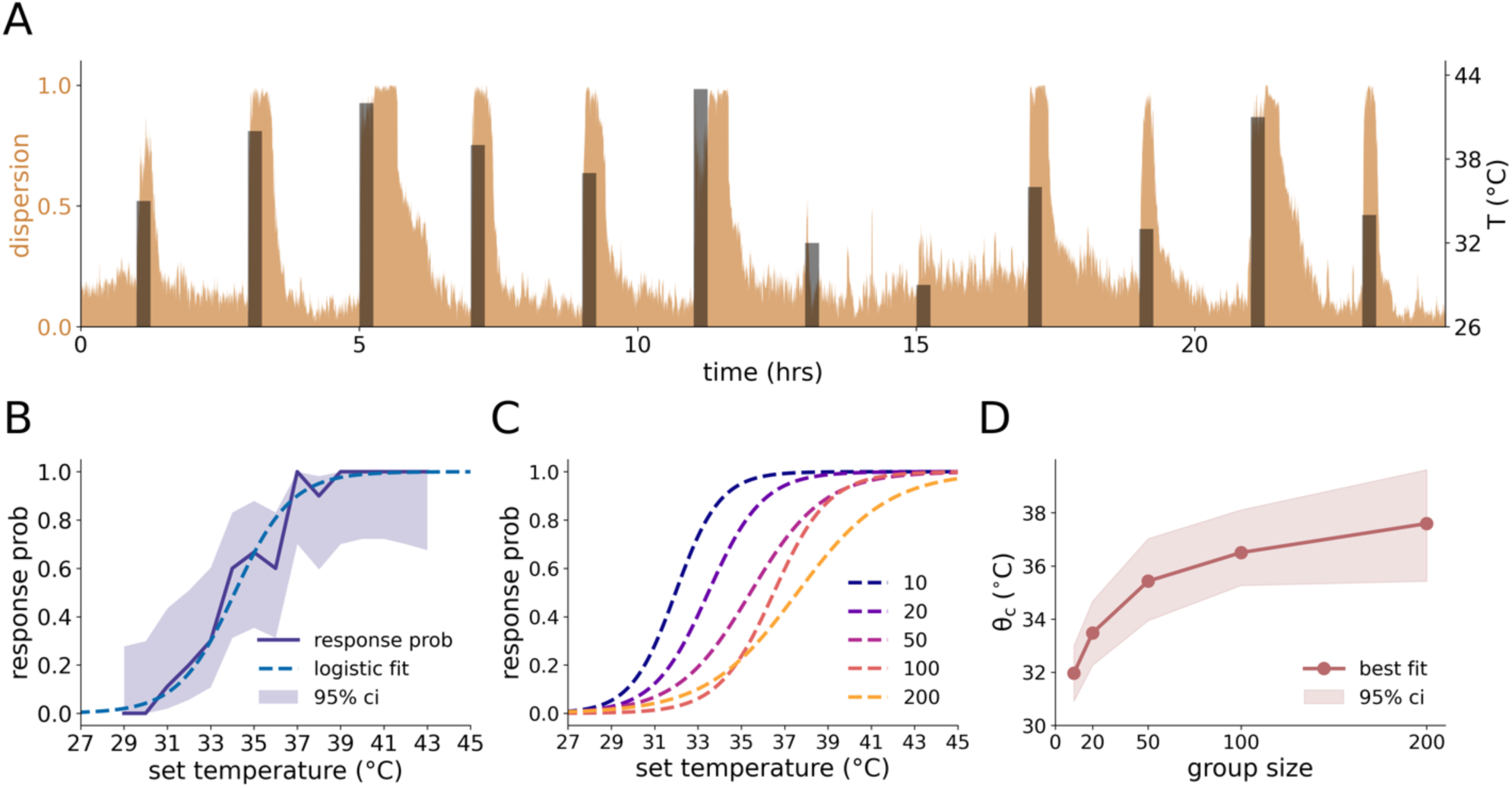
The collective threshold depends on group size. (**A**) A trace from a 24-hour long perturbation protocol using a colony of 36 tagged ants. Perturbations are 15 minutes long and separated by intervals of 2 hours. The set temperature is shown in black. The dispersion of the colony (defined as the fraction of ants outside the nest circle) is shown in brown. The interval allows the colony to relax back to baseline before the next perturbation. (**B**) The probability of a ‘ full response’ (defined as at least 90% of the ants being outside the nest at the same time for at least 30 seconds at some point during the perturbation; solid line) as a function of the perturbation temperature. The light band represents the 95% confidence interval of probability (computed using asymptotic normal approximation for binary coefficient estimation). The dashed line represents a logistic regression fit of the response curve. Experimental colonies consisted of 36 tagged ants. (**C**) Fitted logistic regression curves as in **B** for different colony sizes, showing an upward shift in the response curve. (**D**) The collective threshold parameter *θ*_*c*_, estimated by logistic regression, as a function of colony size. The light band represents the 95% confidence interval, estimated using the bootstrap method with 1,000 sample repetitions (see Methods).

To further investigate this collective threshold, we conducted an experiment with different colony sizes, ranging from 10 to 200 ants. The age, clonal line, and workers to larvae ratio were identical to the experiments described above (Materials and Methods). Each colony was subject to the same experimental protocol with varying temperature perturbations. The collective threshold was estimated for each colony size as above (Figure 3C). Plotting the threshold as a function of colony size (Figure 3D), we found that larger colonies have a significantly higher collective threshold than smaller ones. This effect is robust to variation in the parameters defining the binary collective response (the quorum threshold and duration; Figure S4).

This result is unexpected in light of previously established paradigms for the study of collective sensing, which can roughly be divided into two classes that lead to two different predictions regarding group size effects. The first is ‘ wisdom of the crowd’, in which noisy independent individual estimates of an external signal are pooled to produce a more accurate collective estimate. Under this scenario, the variability in the response is a result of noisy estimation of the external environment by individual ants, and the threshold temperature is an objective quantity that is independent of the group. Accordingly, the prediction would be that larger groups should have higher accuracy (i.e., less variance) in estimating temperature, but the threshold temperature should not change with group size^43–45^. A dependency of the threshold on group size would imply that group dynamics integrate information suboptimally and in a biased way^46– 48^. According to the second paradigm, collective sensing is used to increase the resolution, or sensitivity, to external events such as predator attacks^49,50^. Perturbations of any strength should ideally elicit a response, and the existence of a response threshold is the result of a limited detection capacity. In this case, however, larger groups are predicted to have lower thresholds, because the probability of detecting a weak perturbation increases with the number of individuals in the group. In contrast to these previously documented dynamics, our finding that the collective temperature threshold increases as a function of group size suggests that the response threshold is not an objective quantity to be estimated, but rather a result of a decision-making process that integrates information about the environment with information about the internal state of the colony. It is in fact possible that the optimal collective response threshold differs for colonies of different sizes. For example, if nest evacuation is associated with a relatively higher cost in larger colonies, a threshold that increases with group size might be adaptive.

### The response dynamics are characterized by distinct timescales and social feedback

The strong correlation between the responses of individual ants and the existence of a collective response threshold implies that the collective response is coordinated using interactions between the ants, resulting in positive feedback dynamics. To visualize the dynamics underlying the emergence of the collective response, we plotted the average time course of the collective state (the number of active ants outside the nest, see Materials and Methods) over the perturbation events in the variable-amplitude experiment with colonies of 36 tagged ants (Figure 4A). We divided the perturbation events into three groups, according to the perturbation amplitude: weak perturbations, for temperatures in which the response probability is lower than 0.1; strong perturbations with a response probability greater than 0.9; and intermediate perturbations between those cutoffs. For intermediate perturbations, we separately plotted events in which the colony collective binary response was 1 and 0, respectively. The time course plot is indicative of two dominant processes with distinct timescales. The first is a fast response, with a timescale of 1-3 minutes, in which some ants become excited. This fast response is slower than the timescale of the physical temperature increase (Figure S2E), and the delay might correspond to the internal physiological and neural processing time. At this timescale, the colony response seems to be dominated by the individual responses of ants, and less by interactions between the ants, which leads to a continuous distribution of the colony state variable (Figure 4B). The second process is slower, with a timescale of 5-10 minutes, in which the colony state converges on either a low or a high value (Figure 4B). This convergence indicates the dominance of social feedback, driven by inter-ant interactions during this period.

**Figure 4:**
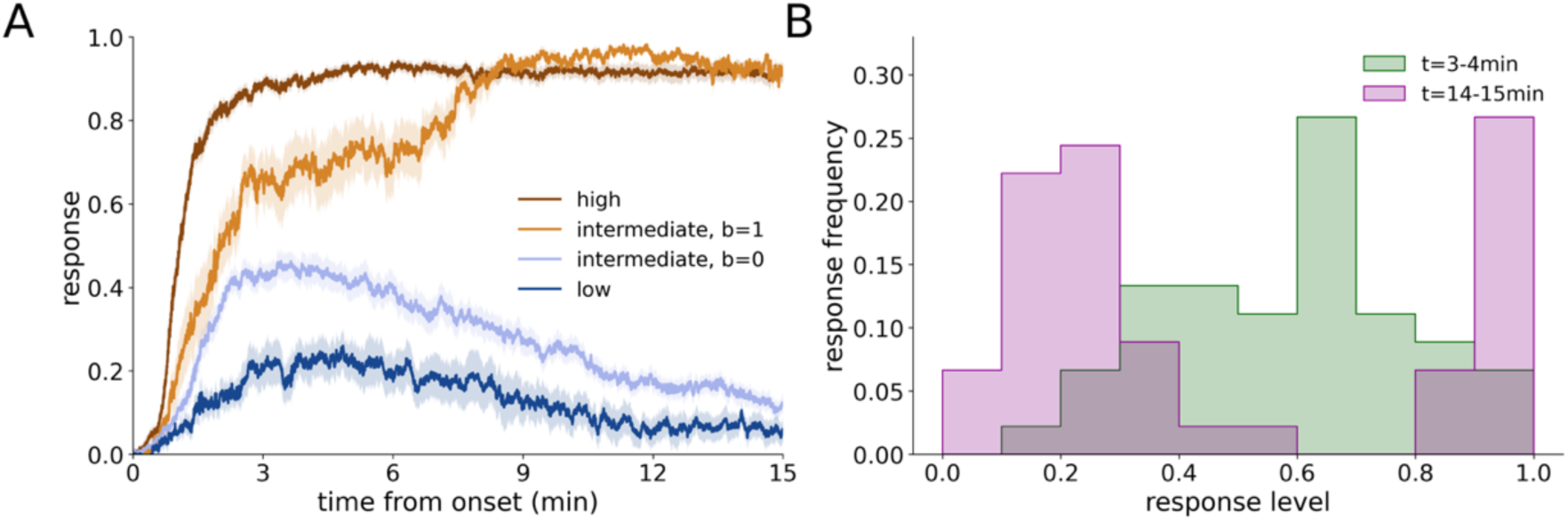
The response dynamics are characterized by distinct timescales and social feedback. (**A**) The evolution in time of the colony state variable (the fraction of active ants) following a temperature change. Single-trial response curves from all experiments with 36 ants were averaged according to temperature and collective response condition. The dark brown curve represents the average response for high temperature, for which the response probability was larger than 0.9. Events without a full response (i.e., b=0) were excluded. The light brown curve represents the average response for intermediate temperature perturbations (response probability between 0.1 and 0.9) in which the colony responded (b=1). Responses in the same temperature range, but in which the colony did not respond (b=0), are depicted by the light blue curve. Finally, the dark blue curve represents the average response for low temperature perturbations (response probability smaller than 0.1; events with positive response (b=1) were excluded). (**B**) Histograms showing distributions of single-trial colony activity states for the intermediate temperature range where the response probability is between 0.1 and 0.9, at two time points along the response curve. The green histogram shows the distribution for the interval between 3 and 4 minutes following perturbation onset, roughly corresponding to a time window in which the effect of the faster process has been exhausted, while the effect of the slower process is not yet apparent. Each datapoint in the histogram is the median value of a single perturbation event in that segment. The pink histogram shows the distribution for the interval between 14 and 15 minutes, at which both transient dynamics of the response have run their course.

### A binary network model recapitulates the emergence of a collective threshold

To better understand the interactions that might underlie a colony’ s collective response, we implemented a simple spin-like binary network model. This type of model is commonly used for complex and collective systems^51–54^. We represent each ant by a binary variable *σ*_*i*_, in which *σ*_*i*_ = 1represents the ‘ perturbed’ behavioral state, and *σ*_*i*_ = −1 the ‘ relaxed’ behavioral state (Figure 5A). The behavioral state of each ant is decided by a logistic activation function (Figure 5B):

**Figure 5:**
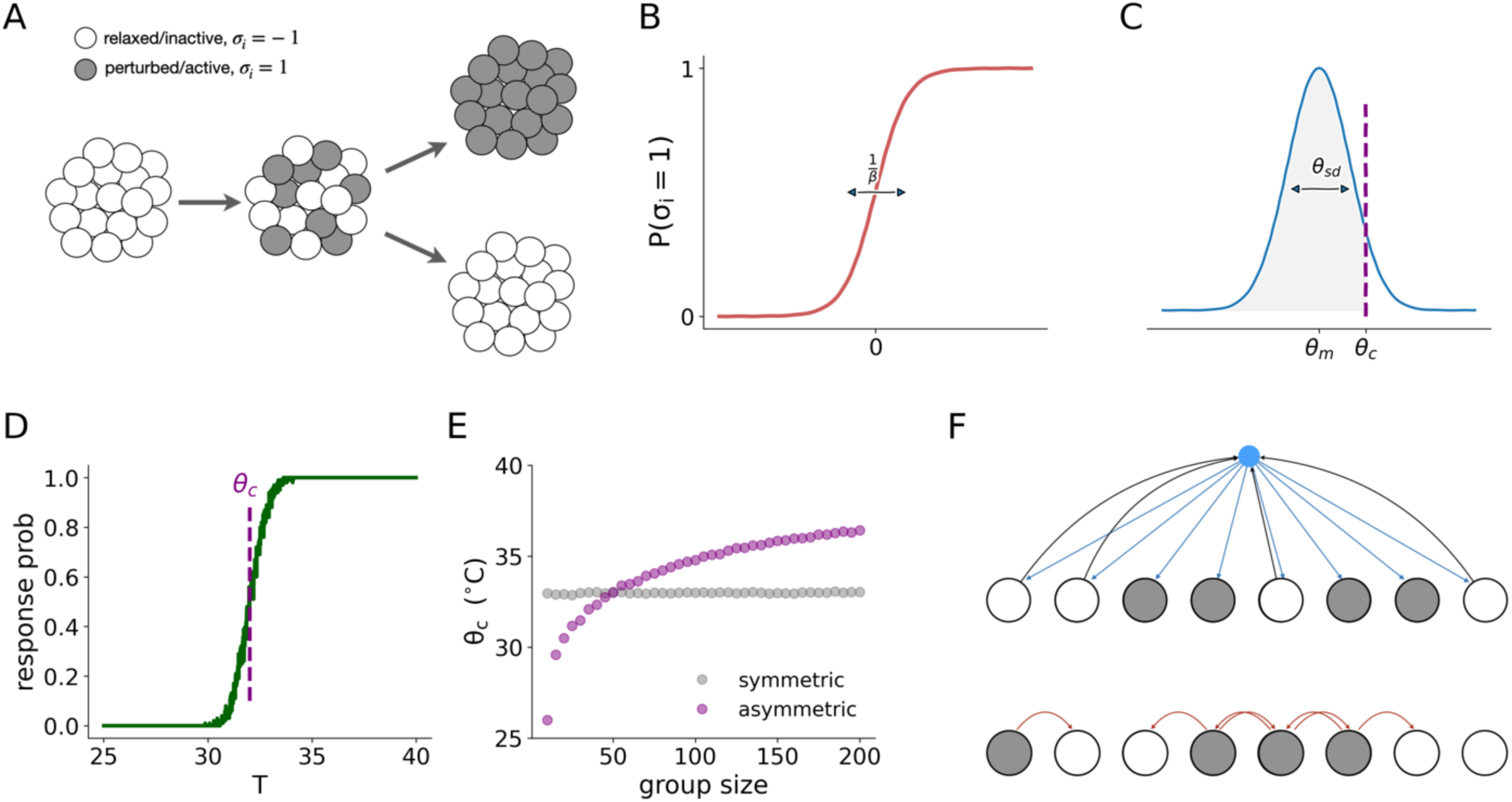
The emergence of the collective threshold can be modelled with two opposing forces. (**A**) An illustration of the model’ s two-phase dynamics. In the first phase, the ants respond independently according to their individual response thresholds. As a result, a subset of ants become active. In the second phase, the interactions between the ants result in the colony being either fully active or fully inactive. (**B**) The logistic activation function of the individual ant. The ant is either active or inactive in a probabilistic manner depending on an integrated input parameter *h*_*i*_. A thermal noise parameter *β* controls the width of the ambiguous response region. (**C**) The individual response thresholds *θ*_*i*_ are sampled from a normal distribution of mean *θ*_*m*_ and width *θ*_*sd*_. The collective threshold *θ*_*c*_ is the temperature for which the cumulative probability equals *m*_*c*_. (**D**) Simulation of the collective threshold, showing the response probability as a function of temperature, averaged over 100 simulation runs. For each run, a new set of individual thresholds is sampled. See Materials and Methods for full details on the simulation parameters. (**E**) The collective threshold as a function of group size for the basic model (grey circles) and the asymmetric model (purple circles). The interaction parameters *j*^*p*^ and *j*^*r*^ were chosen to have the same collective threshold at *N* = 50 and to approximately replicate the range of thresholds observed in the experiment. (**F**) An illustration of possible ant interaction mechanisms. Global, pheromone-based interaction (top), in which ants in a given state contribute (black arrows) to the total concentration of pheromone in the environment (blue circle), which is then perceived by all ants (blue arrows). Local, contact-based interaction (bottom), in which ants in a given state only affect the behavior of nearby ants (red arrows).

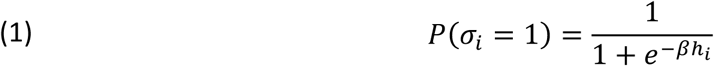

Here, *β* is the thermal noise parameter, and *h*_*i*_ is the integrated input of ant *i*. In the general case, this integrated input is a combination of the external temperature perceived by the individual ant and the contribution of the social interactions. For simplicity, we take advantage of the apparent separation between the timescales of these two components and assume the dynamics to have two phases (Figure 5A). In the first phase, the ants respond individually and independently to the external temperature, according to the equation:

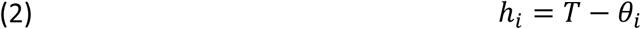

Here, *T* is the perturbation temperature and *θ*_*i*_ is the individual threshold of ant *i*, randomly drawn from a normal distribution with mean *θ*_*m*_ and standard deviation *θ*_*sd*_ (Figure 5C). Following this phase, some of the ants will be in the relaxed state and some in the perturbed state (Figure 5A, middle). The state of the colony is characterized by the order parameter *m*, defined as the fraction of perturbed ants in a colony with *N* ants:

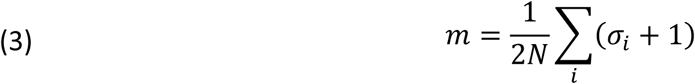

In the second phase, the states of the ants change over time according to the social dynamics:

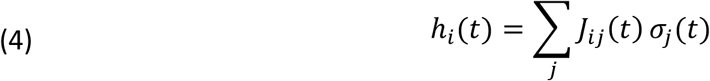

Here, *j*_*ij*_(*t*) is the interaction strength between ant *i* and ant *j* at time *t*. In the most general case, the interaction between a pair of ants will depend on their relative position, and on their behavioral states. We further simplify our model by ignoring space and assuming that all ants interact with all other ants all the time. This can be justified by the observation that the collective response dynamics of the ants are much slower than their movement speed in the nest during the ‘ excitement’ period that precedes the response (Video 1). Under this assumption, we can characterize the interaction by two parameters, representing the exciting effect of perturbed ants and the inhibiting effect of relaxed ants. The input to all the ants is then the same, and can be written as:

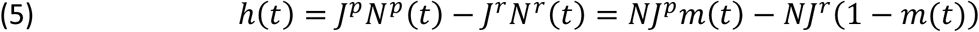

Where *j*^*p*^ and *j*^*r*^ are the interaction parameters, and *N*^*p*^ (*t*) and *N*^*r*^ (*t*) are the number of perturbed and relaxed ants at time *t*, respectively.

Assuming the thermal noise is low enough, the model is characterized by two stable fixed points at *m* = 0 and *m* = 1, representing the collective relaxed and perturbed states (Figure S5A-D), with a separatrix at:

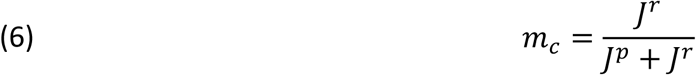

This implies that, on average, the value of *m* at time zero will decide the final collective state of the colony. This *m*_0_ is a result of the individual responses of the ants to the temperature increase during the first phase of the dynamics. Therefore, the threshold perturbation temperature *θ*_*c*_ is the temperature at which the average fraction of perturbed ants will equal *m*_*c*_. In the case where the thermal noise is small compared to the individual threshold variability, this threshold satisfies the condition:

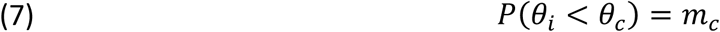

The exact value of this threshold will depend on the distribution of the individual thermal thresholds in the ant population, as well as the ratio between the excitatory and inhibitory interaction strengths (Figure 5C; Figure S5).

### Group size dependency of the collective threshold entails asymmetric interactions

While the model, as defined above, captures the emergence of the collective threshold from the interactions between the ants (Figure 5D), it does not include any group size dependency that can reproduce the experimental observations in Figure 3 (Figure S5L; grey curve in Figure 5E). Because ants in the experiments had a narrow age range, belonged to the same clonal line, and were sampled randomly from the same stock colony in each experiment (Materials and Methods), we can assume that their individual thermal thresholds were also sampled from the same distribution, regardless of colony size. This implies that, under the assumptions of the model, group size dependency can only arise if the excitatory and inhibitory components scale differently with group size *N*. For example, we can set the inhibitory interaction to scale linearly with *N*, but let the excitatory interaction be independent of *N*:

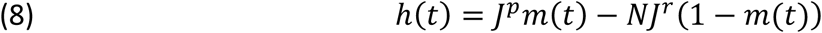

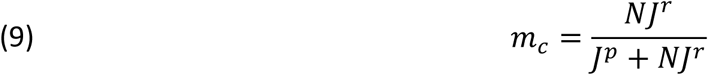

Using these definitions, the threshold value will sub-linearly increase with the size of the colony (purple curve in Figure 5E).

This choice is not arbitrary, because different types of interactions should produce different scaling with *N*. Clonal raider ants are blind, and mostly communicate via pheromones and tactile interactions. For a volatile pheromone, the concentration in the air surrounding ants aggregated in close proximity in the nest should scale with the number of ants that emit the pheromone, which equals the fraction of pheromone emitting ants multiplied by *N*. This pheromone concentration is then perceived by all ants in the colony (Figure 5F top). The exact form of the scaling will depend on the chemical properties of the pheromone, the physical properties of the environment, and the concentration/response curve of the ants themselves. On the other hand, physical interactions between pairs of ants will depend on the rate of encounters. Physical interaction is a common excitatory mechanism in ants, particularly in scenarios of recruitment ^55,56^. When the colony is dense, the encounter rate per ant saturates, meaning an ant is always in contact with other ants at the maximum capacity. This implies that the total excitatory force an ant feels is dependent on the fraction of active ants in the colony and does not scale with the size of the colony (Figure 5F bottom). Of course, the distinction between the two mechanisms does not have to be clear-cut, and can be quantified with a scaling parameter *α*:

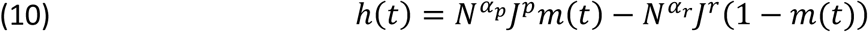

The value of *α* represents a gradual transition between local, nearest-neighbor interactions and global, all-to-all interactions. An increase of collective threshold with group size will then emerge for any *α*_*r*_ > *α*_*p*_ (Figure S6).

So far, we have considered interactions that are state-dependent, that is, interactions in which ants signal their state to other ants. However, some interactions within the colony can be regarded as state-independent, or to vary slower than the typical timescale of the behavioral response. For example, weakly volatile “aggregation” pheromones could mark the nest site. The strength of this nest odor will then scale with the number of ants in the nest but will not change because of ants leaving the nest momentarily. The existence of such a pheromone is supported by the tendency of the ants to settle back at their original nest location following perturbations (Figure S7). Such an aggregation signal would act as a constant pulling force that balances the excitation within the colony. In our model, we can account for such an interaction by removing the state dependency from the inhibitory term, and write equation (10) as:

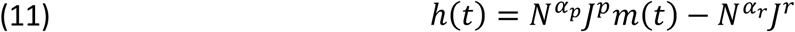

Because the scaling of the inhibitory interaction strength is the same as in the initial version of the model, we again get an increase of the threshold with group size (Figure S6D-E). However, the shape of the increase and the predictions of the model for larger group sizes differ (Figure S6F).

## Discussion

Our results provide a simple, tractable example of a collective perception-action loop, where social dynamics are used to integrate the sensory perception of individual ants and to produce a coherent collective response. We show that under borderline conditions, individual ants suppress their own assessment or perception of sensory information about the external environment in favor of a collective decision. Moreover, the social dynamics enable the colony to integrate information not only about the external environment, but also about the state of the colony itself (its size in this case). The collective outcome is therefore more than a mere average of the “opinions” of the individual ants.

The collective sensory threshold seems to emerge from a balance between two opposing forces. As in other biological systems, these forces do not have to map to a single biological mechanism, but could rather represent a combination of various processes with a similar functional effect. For example, excitation and inhibition in a neural network arise from many different types of neurons and neurotransmitters whose effects differ in aspects such as timescale, spatial distribution and plasticity. Likewise, transmembrane currents are the product of many types of ion channels, each modulating the membrane response in a different way. This mechanistic complexity underlies both the robustness of biological systems and their flexibility to adapt their responses to various conditions on multiple timescales^57–62^. Similarly, the excitatory and inhibitory forces at play in an ant colony are likely composed of various chemical, physical, and possibly other types of interactions. The isolation of individual mechanisms and the understanding of their precise functional role in the collective dynamics will require further experiments. However, mesoscopic models such as the one employed here can provide a formal understanding of the principles of emergent collective computation even without detailed knowledge of the underlying mechanisms.

## Materials and Methods

### Experimental setup

The 10×10 cm behavioral arena consists of a 3mm thick plaster of Paris layer on top of a temperature-controlled metal platform. The platform is divided into 4 zones. The temperature of each zone is measured by an embedded thermistor (Omega 44304) and controlled by a thermoelectric cooling (TEC) device (CUI Inc. CP60). The plaster arena is surrounded by a metal frame heated to a high temperature (between 45°C and 50°C) to confine the ants. The arena and the frame are installed inside a box in which the air temperature and the relative humidity (RH) are continuously monitored. Figure S1 gives full details and a schematic of the experimental setup.

### Temperature control

The temperature in each of the arena zones is regulated by an Arduino-implemented PID controller. An additional PID is used to control the temperature of the heated frame. The spatial integrity of the surface temperature is verified by an IR thermal camera (FLIR Lepton 3.0). Details of implementation and verification are given in Figure S2. All temperature values reported are the set temperature of the PID, while actual ground temperatures can be inferred from the calibration curve (Figure S2B).

### Moisture control

*O. biroi* ants are sensitive to moisture and humidity, and a drop in these parameters will affect their behavior. Without additional measures, heating the plaster floor during temperature perturbations would lead to desiccation. We therefore developed a method to maintain a stable moisture level of the plaster during the experiments. We found that the color of the plaster is a robust and reliable measure for moisture level (the plaster gets lighter/brighter as it dries). We therefore defined the ‘ moisture index’ as the 90^th^ percentile of the pixel brightness value distribution in the arena (Figure S3A-B). This definition is robust to the presence of ants and accumulation of trash in the arena during the experiment, because both contribute to the low-brightness tail of the distribution. As the baseline color of the arena depends on batch effects of the plaster, as well as on the exact tuning of the camera and illumination, the set point was manually determined before each experiment. At the start of each experiment, we calculated the pixel brightness distribution and the moisture index for the arena when dry and when completely saturated with water (Figure S3B). We then defined the threshold moisture index *M*_*c*_ as:

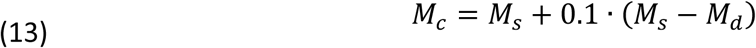

*M*_*s*_ and *M*_*d*_ are the respective indices under saturated and dry conditions. During an experiment, the moisture index was calculated every second (Figure S3C), and whenever it rose above threshold, the waterflow into the embedded water delivery tubes (Figure S1) was opened for a short (0.5 sec) duration. Figure S3C depicts an example time course of the moisture index when the control is enabled (left of the dashed line), and when it is disabled, and the plaster is allowed to dry (right of the dashed line).

### Experimental design

Experimental colonies were composed of age matched, one cycle old workers, and 6-7 days old larvae in a 2:1 worker to larvae ratio. All ants were derived from the same stock colony (STC6), which belongs to *O. biroi* clonal line B^63^. For the threshold experiment, two data sets were collected: one with individually tagged ants in 10 colonies of 36 adults and 18 larvae each, and one with untagged ants and variable colony sizes of 10, 20, 50, 100, and 200 ants, as well as 5, 10, 25, 50, and 100 larvae, respectively. For the latter data set, each group size was represented by 3 replicate colonies (15 colonies total). Colonies were assayed sequentially, one per week, between October 2018 and August 2019. The experiments did not have any specific order. For the collectivity measurement experiment, 4 colonies of 36 tagged ants and 18 larvae were used.

### Colony preparation

Five day old workers were separated every other week from an *O. biroi* stock colony and split into two experimental colonies. These ants laid eggs ca. 10 days later. In the first colony, we waited for larvae to hatch, and began experiments when larvae were 6-7 days old. In the second colony, we removed the first batch of eggs after 3 days. This resets the colony’ s reproductive cycle and creates a one-week developmental lag between the first and second colony, allowing us to run experiments continuously. Right before experiments began, colonies were adjusted to the number of ants and larvae required for the experiment of that week and transferred into the experimental setup.

### Color tagging

For experiments with individually tagged ants, 12 day old ants were marked with color dots on the thorax and gaster using oil-paint markers (UniPaint Markers PX-20 and PX-21)^21,64^. Colonies for these experiments contained 36 ants, marked with all unique combinations of blue, green, orange, pink, purple/red and yellow.

### Experimental protocol

After experimental colonies had been transferred to the experimental setup, they were allowed to settle for 48 hours. During that time, the set temperature of the arena was 26°C and the ants were fed fire ant (*Solenopsis invicta*) pupae. Six hours after the last feeding event we began the perturbation protocol, and we did not feed the ants for the rest of the experiment. Every 2 hours, the set temperature of the arena was increased to the perturbation value for 15 minutes, and then lowered back to the baseline temperature. For threshold measurements, the sequence of perturbations was a random permutation of the values between 29°C and 43°C. The sequence was presented in the same order for all colonies in the experiment. For the collectivity experiment, colonies were subjected to a sequence of identical perturbations of either 40°C or 33°C.

### Video recording and tracking

Videos were recorded at 10 frames per second using a FLIR Flea3 camera (image size 2500×2500 pixels). The videos were analyzed using anTraX, a software package for video tracking of color tagged ants^42^. For experiments with tagged ants, the software outputs an estimated location for each ant in the colony in each frame. For experiments with untagged ants, the software outputs a list of segmented blobs containing ants, together with their respective centroid coordinates and area.

### Defining the location and size of the nest

Most of the behavioral measures we use in our analysis depend on the location of the nest. Because we do not use a physical nest structure, and because the nest location is dynamic and changes during an experiment, we estimate the location and size of the nest from the tracking data. Typically, 50-90% of the ants in the colony will reside in the nest at any given time. Therefore, the median location of all the ants will give an accurate estimate of the nest centroid. To be consistent between tagged and untagged experiments, we do not use the individual location data for estimating the nest location. Rather, we use the relative area of each blob as a proxy for the number of ants in that blob, and the equation to determine the nest location takes the form:

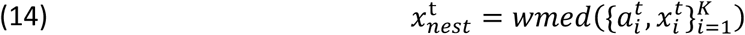

In this equation, 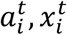 are the area and the coordinates of the *i*-th blob at time *t, K* is the number of blobs in the frame, and *wmed* is the weighted median function. We further use a median filter with duration *k* = 10 *min* to suppress fluctuations in this measure:

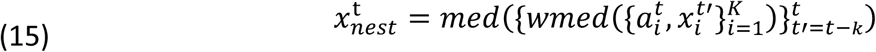

We chose the nest to have an effective radius *R*_*nset*_ equal to the major semi axis of the nest blob. The nest blob, as segmented by the tracking algorithm, represents the tight aggregation of ants in the center of the nest. As in the case of the nest location, we use median filtering to smooth fluctuations. For the analysis of individual and collective responses to perturbations, we use the nest location and radius at the onset time of the perturbation.

### Individual response measures and collectivity

We define an individual ant as having responded to the perturbation when she has first exited a circle of radius 15mm around the nest for a duration of at least 30 consecutive seconds (Figure 2A). This radius was chosen so that ants that are excited or active near the nest will be mostly inside the circle, while ants evacuating the nest will exit the circle. We then characterize the response of each ant for a given perturbation using three measures. The binary response 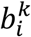 is set to 1 if the *i*-th ant leaves the circle during the duration of the *k*-th perturbation for a consecutive period of at least 30 seconds. The response delay 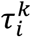 is defined as the delay between the onset of the perturbation to the time the ant first crossed the circle for such a period, and the response direction 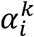 is defined as the angle of the point of crossing from the line connecting the nest’ s centroid to the center of the arena. We then calculate the correlation between the measures for each pair of ants in the same colony. In case an ant was outside the circle at the onset of the perturbation, we treat its response to that particular perturbation event as a missing value. We generate a null distribution for these measures by randomly permuting the responses of each ant to all the perturbations in the experiment, and calculating the distribution of pairwise correlation coefficients for this shuffled dataset.

### Measuring the collective threshold

We define the colony activity variable 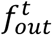 as the fraction of ants in the colony that are outside the nest at time *t*. As before, to be consistent across experiments with tagged and untagged ants, we do not use the individual location data, but rather an estimate based on blob sizes:

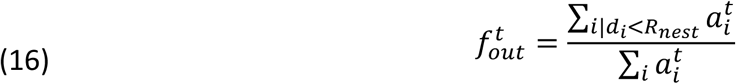

Here, *d*_*i*_ is the distance of the *i*-th blob from the nest, and *R*_*nest*_ is the effective nest radius at the beginning of the perturbation, as defined above. Note that in the context of the collective state, we use a smaller circle around the nest than in the case of the individual response, in order to better capture the activity of ants around the nest, and not only ants evacuating the nest.

We define the colony as having ‘ responded’ if 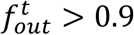 for a period of at least 30 consecutive seconds during a perturbation. To estimate the response threshold, the responses of 9 colonies in the experiment with tagged ants were pooled. Events in which the ants were not well settled at the beginning of the perturbation were excluded from the analysis. For inclusion, we required the average number of active ants in the 15 minutes preceding the onset of the perturbation to be lower than half the median value, calculated across all the events in the experiment. This resulted in the exclusion of 8 of the 150 events. The threshold was estimated by fitting a logistic regression model to the collective binary response variable. Confidence intervals for the threshold were estimated using the case resample bootstrap method with 1000 replicates. For the group size experiment, we repeated the analysis above, resulting in a threshold parameter and confidence interval for each group size (Figure 3D). The effect of colony size was estimated using a logistic regression model with group size and temperature as the independent variables (Figure S4).

### Simulation of the mathematical model

The binary network model was simulated using the asynchronous update approach. In each simulation step a randomly selected ant updates its state. In each simulation run, the ants were initialized at the inactive state *σ*_*i*_ = −1. Each run lasted 20 full update cycles; an update cycle is defined as *N* simulation steps, where *N* is the number of ants in the simulated colony. The full simulation parameters are given in the figure captions.

All parameter sweeps were performed on the Rockefeller University high performance computing cluster.

## Acknowledgements

We thank J. Saragosti of the Laboratory of Social Evolution and Behavior and J. Petrillo of the Rockefeller University Precision Fabrication Facility for help in designing, building and troubleshooting the experimental setup. L. Olivos Cisneros and S. Valdés Rodríguez helped setting up experimental colonies. We also thank S. Leibler, G. Maimon and A. J. Libchaber for fruitful discussion and advice. Research reported in this publication was supported by the National Institute of General Medical Sciences of the National Institutes of Health under Award Number R35GM127007 to D.J.C.K. The content is solely the responsibility of the authors and does not necessarily represent the official views of the National Institutes of Health. This work was also supported by a Searle Scholar Award, a Klingenstein-Simons Fellowship Award in the Neurosciences, a Pew Biomedical Scholar Award, and a Faculty Scholar Award from the Howard Hughes Medical Institute to D.J.C.K. A.G. was supported by the Human Frontiers Science Program (LT001049/2015). This is Clonal Raider Ant Project paper #19.

---

**Figure S1:**
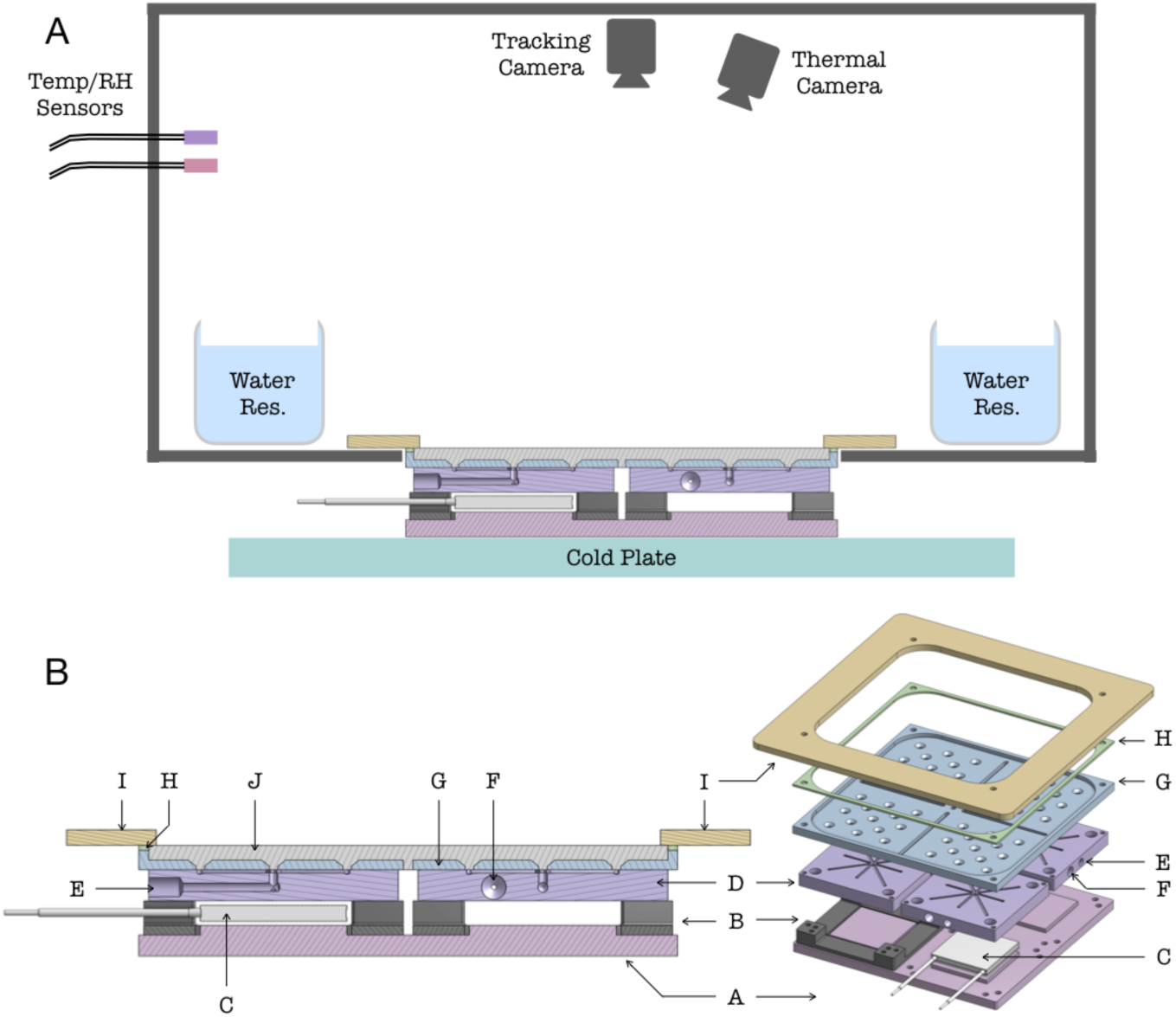
Experimental setup. (**A**) Schematic of the experimental setup (not to scale). The plaster arena and the temperature control platform are situated inside a closed box, together with a water reservoir to reduce the rate of humidity loss from the plaster. The ambient temperature and relative humidity are continuously monitored. A tracking camera records movies of the ants’ behavior, and a thermal camera monitors the temperature on the arena’ s surface. The temperature control platform is placed on a cold plate to sink the excessive heat generated by the thermoelectric coolers (TECs). The custom electronic controls used to drive the TECs and the water flow, and the computer with a custom software used to collect and record data (movies and synchronized temperature and humidity measurements), are not shown. (**B**) Detailed to-scale schematics of the temperature control platform. A: Bottom metal base layer. B: Thermally insulating spacer and support, one for each TEC device (not all are shown). C: TEC device, one for each zone. Thermally conductive silicone pads that attach the device to the metal parts below and above are not shown. D: Middle metal layer, which contains the water tubing and the zone’ s thermistor. This layer is made of 4 pieces separated from each other to enable individual control of each zone. E: Tubing for water flow, machined into the middle metal piece. Each metal piece contains one inlet which opens up at the surface to distribute the water to 13 outlets into the plaster. F: Hole for thermistor. G: Top metal plate, in which the plaster arena is cast. The top plate is divided into 4 separate zones, which are connected with minimally sized bridges to enable separate temperature control of the four zones (this feature of the setup was not used in the current study; all zones were set to the same temperature in all experiments). The top metal plate is attached to the middle metal pieces (D) using thermally conductive silicone pads (not shown). H: Thermal insulation layer between heated barrier and top metal layer. I: Heated metal barrier (3mm thick aluminum plate). Not shown are the attached thermistor and heating resistors used to control the barrier’ s temperature. J: Plaster arena, cast onto the top metal plate (G).

**Figure S2:**
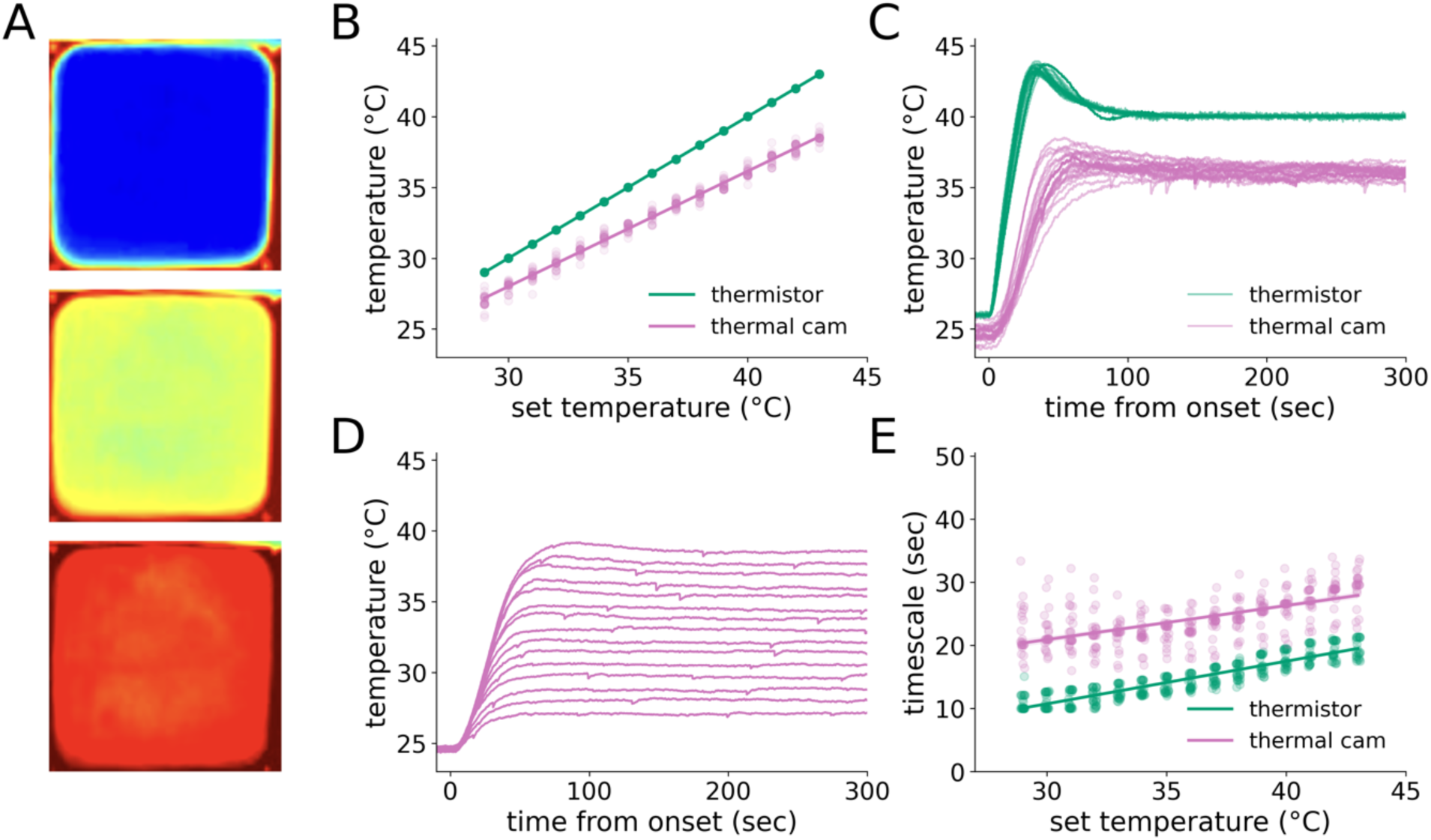
Temperature control. (**A**) Snapshots from the thermal camera during an experiment, for set temperatures of 26°C, 34°C and 40°C (top to bottom). The temperature-controlled arena is surrounded by a high-temperature heat fence which is used to confine the ants to the arena even under the maximum perturbation. (**B**) Calibration curve showing the actual temperature vs. the set temperature. The plot shows the thermistor measurement (green), and the temperature measured by the thermal camera (pink). Each scatter point is data from one perturbation, and the plot shows pooled data from all experiments analyzed in this paper. Both the thermistor and thermal camera measurements are time averages of the measurements between 5 minutes and 15 minutes after the onset of the perturbation. The thermistor measurements are averaged over the 4 thermistors of the arena, and the thermal camera measurements are averaged over the pixels of the arena region of interest. Because the thermistor measurements provide feedback to the TEC controllers, they display 1:1 correspondence with the set temperature with minimal variability. The thermal camera curve is linear with a slope of slightly less than 1, implying a temperature drop across the layer of plaster. The variability in the y-axis is mostly due to the instability in the thermal camera measurements between experiments, and not because of actual variability in temperature across experiments. Because of the low reliability of the thermal camera absolute measurements, we do not use them for control, but only for analysis of the spatiotemporal dynamics of the arena’ s temperature. (**C**) Comparison of the temperature dynamics as measured by the embedded thermistor (used for temperature control, green) and the thermal camera (pink), for a perturbation setpoint of 40°C. While the temperature underneath the plaster climbs fast and displays a significant overshoot as a result of the PI control algorithm, the ground temperature changes slower and the overshoot is dampened by the mass of the plaster layer. (**D**) Temperature traces from the thermal camera, averaged across perturbations with the same set temperature, for set temperatures between 29°C and 43°C. (**E**) The rising timescale of the single-perturbation temperature trace, defined as the latency from the setpoint change time to the half rise time of the temperature, for the thermistor and the thermal camera. The timescale was estimated by fitting a single exponential to the trace between 10 seconds and 60 seconds following the onset of the perturbation. There is a slight increase in timescale for higher temperatures. This increase is expected because of the use of a PI controller to set the TEC current. The range of variation in the heating timescale is much lower than the typical response time of the colony (Figure 4A).

**Figure S3:**
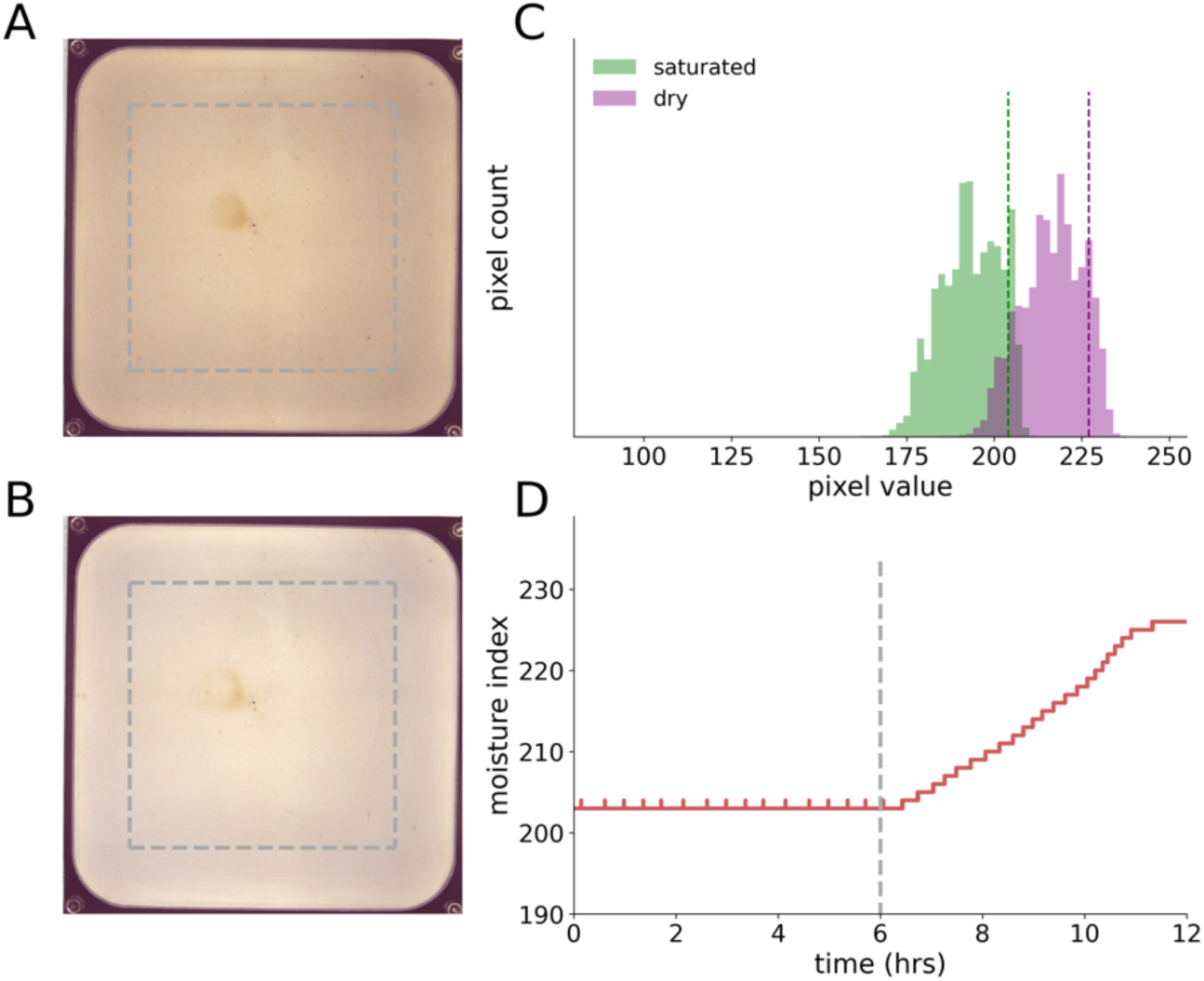
Moisture control. (**A-B**) To calculate the moisture index, the distribution of pixel brightness levels in the arena, excluding margins of 1cm width (dashed grey line), is calculated. Depicted are examples of a fully saturated (**A**) and a completely dry (**B**) frame. (**C**) At the start of each experiment, the pixel brightness distribution is calculated for the arena when dry (purple), and when completely saturated with water (green). A moisture index is then calculated from each distribution as the 90^th^ percentile of the brightest pixels (dashed lines). (**D**) An example time course of the moisture index when moisture control is enabled (left of the dashed line), and when it is disabled, and the plaster is allowed to dry (right of the dashed line). The small spikes in the left part of the curve represent times at which the index increases above threshold and is immediately compensated by the controller.

**Figure S4:**
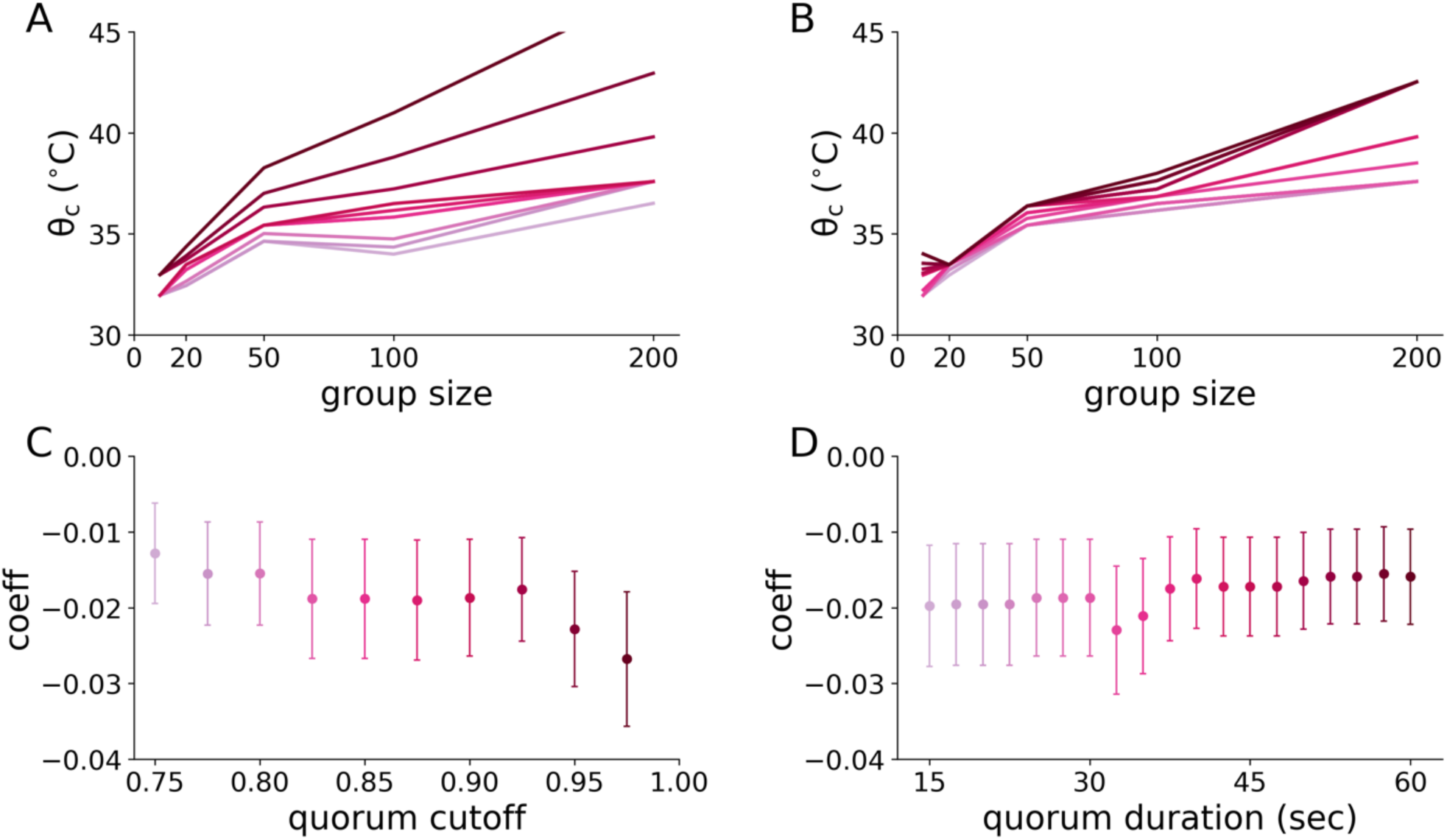
The effect of group size is robust to changes in analysis parameters. (**A**) The threshold as a function of group size for different values of the quorum cutoff parameter, from 0.75 (light) to 0.975 (dark). The threshold data were generated as in Figure 3D. (**B**) The threshold as a function of group size for different values of the quorum duration parameter, from 15 seconds (light) to 60 seconds (dark). The threshold data were generated as in Figure 3D. (**C-D**) The effect of group size on response probability is significant for all tested parameter combinations. The figure depicts the coefficient of the group size parameter in the logistic regression model, together with its 95% confidence interval, for each parameter value tested. Negative coefficients imply a decrease in response probability (i.e., an increase in threshold) with group size.

**Figure S5:**
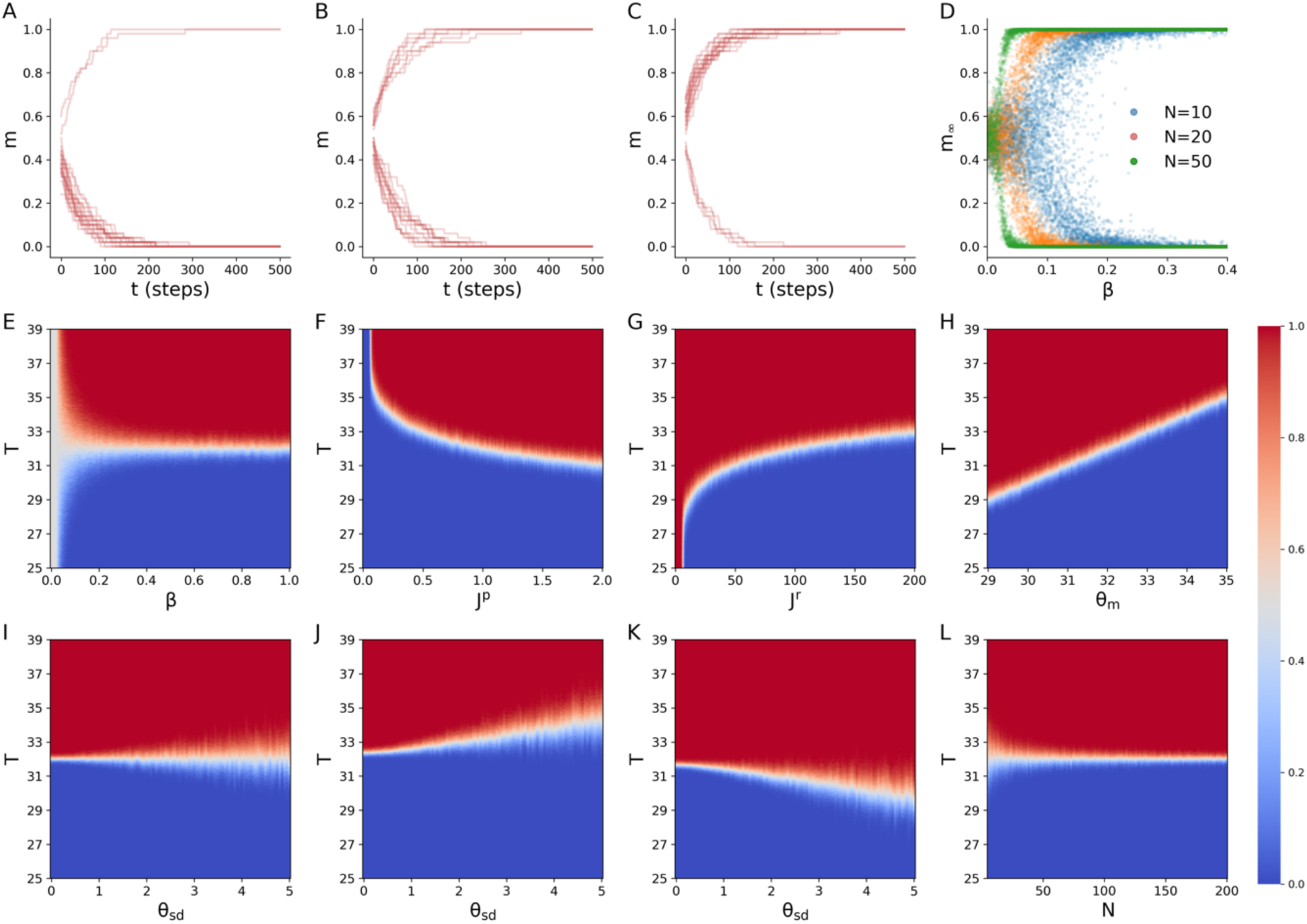
Exploration of the basic model. Unless otherwise stated, all simulation runs are performed for a duration of 20 update cycles. Each cycle consists of *N* asynchronous updates, with *N* being the number of ants in the colony. Default parameter settings are = 50, *β* = 1, *j*^*p*^= *j*^*r*^ = 1, *θ*_*m*_ = 32, *θ*_*sd*_ = 2. For all heatmaps, each pixel represents the average collective response across 100 independent runs of the model. (**A-C**) Evolution of the colony state *m* as a function of time (measured in cycles), for temperature perturbations of 31.5 (A), 32 (B) and 32.5 (C). For each condition, 25 runs are shown. For threshold temperature perturbations (B), the probability of a colony to respond with a collective nest evacuation is 50%, and the outcome in each case depends on the specific values of individual thresholds of the ants, as well as on the thermal noise. For temperatures below (A) or above (C) the threshold, the probability of a collective response is smaller or larger than 50%, respectively. (**D**) The collective threshold (i.e., the existence of two fixed points for high and low *m* with a separatrix between them) is contingent on sufficiently low thermal noise. The plot shows the final *m* for *β* in the range 0-0.4, with 25 runs of each value of *β*, for 3 colony sizes. Smaller colonies are more sensitive to thermal noise, and require a larger value of *β* in order to have a collective threshold. (**E**) The value of the collective threshold does not depend on the thermal noise. The plot shows a heatmap of the collective response probability as a function of temperature and *β*. Subsequent simulations are therefore done at a high *β* so that thermal noise is negligible for all colony sizes. (**F-G**) The effect of the interaction parameters *j*^*p*^ (F) and *j*^*r*^ (G) on the collective threshold. As predicted from equation 6, the threshold decreases with *j*^*p*^ and increases with *j*^*r*^. (**H**) The collective threshold directly corresponds to the individual threshold average *θ*_*m*_. When *j*^*p*^= *j*^*r*^, the collective threshold equals *θ*_*m*_. (**I-K**) The effect of the individual threshold variability *θ*_*sd*_ on the collective threshold is contingent on the ratio of *j*^*p*^and *j*^*r*^. If *j*^*p*^ = *j*^*r*^ (I), there is no effect on the collective threshold. If *j*^*p*^ < *j*^*r*^ (J), then *m*_*th*_ > 0.5, and as the distribution of individual thresholds gets wider, the collective threshold is increasing. If *j*^*p*^ > *j*^*r*^ (K), then *m*_*th*_ < 0.5, and as the distribution of individual thresholds gets wider, the collective threshold is decreasing. (**L**) The collective threshold in the basic model does not depend on colony size *N*. This holds regardless of the value of the other parameters in the model.

**Figure S6:**
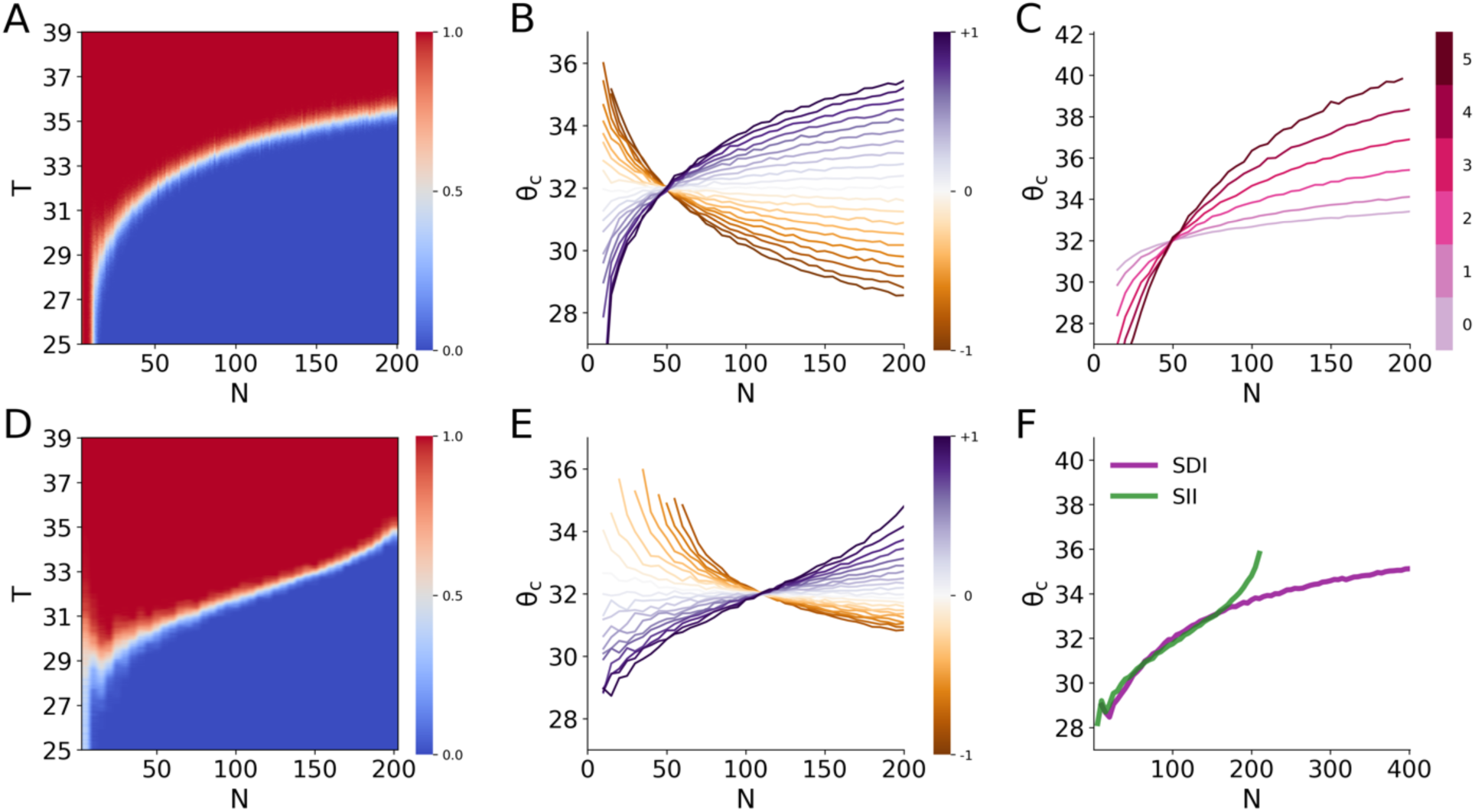
Model with interaction asymmetry. Unless otherwise stated, all simulation runs are performed for a duration of 20 update cycles. Each cycle consists of *N* asynchronous updates, with *N* being the number of ants in the colony. Default parameter settings are = 50, *β* = 1, *j*^*p*^ = *j*^*r*^ = 1, *θ*_*m*_ = 32, *θ*_*sd*_= 2, *α*_*r*_ = 1, *α*_*p*_= 0. For heatmaps, each pixel represents the average collective response across 100 independent runs of the model. (**A**) Heatmap of the collective response probability as a function of temperature and group size, as in Figure S5L, but with the asymmetric interaction of Equation 8. (**B**) The collective threshold as a function of group size for different value combinations of the scaling parameters *α*_”_ and *α*_*i*_. For each curve, the parameter values were set to *α*_*r*_ = 1+ *α* and to *α*_*p*_ = 1− *α*, where the value of *α* is ranging from -1 (decreasing threshold, brown) to 1 (increasing threshold, purple). The interaction parameters *j*^*p*^and *j*^*r*^ were scaled so the inhibitory and excitatory components are equal for *N* = 50, resulting in a collective threshold that equals the mean individual threshold. (**C**) The collective threshold as a function of group size for *α*_*r*_ = 1, *α*_*p*_ = 0, but with different values for *θ*_*sd*_, showing that the variability in individual thresholds determines the magnitude of change in the collective threshold with group size. (**D**) Heatmap of the collective response probability as a function of temperature and group size, as in **A**, for the state-independent inhibition model (Equation 11). (**E**) The collective threshold as a function of group size for the state-independent inhibition model (Equation 11) for different value combinations of the scaling parameters *α*_*r*_ and *α*_*p*_ (values are related, and curves are color coded as in **B**). (**F**) The state-dependent and state-independent inhibition models extrapolate differently to larger colony sizes, giving different predictions. For the state-dependent inhibition (SDI, purple), the collective threshold increase slows down as colony size increases. This is because no matter how strong the inhibition, once the temperature is above the highest individual threshold in the colony, it zeroes out after the first phase of the dynamics. For the state-independent inhibition (SII, green), the collective threshold increase accelerates as colony size increases, until at a critical colony size the colony becomes unresponsive regardless of the temperature value. This is because the inhibition becomes so strong that even if all individuals are perturbed in the first phase, this is still not sufficient to overcome the inhibition, and the ants eventually relax back. The interaction parameters *j*^*p*^ and *j*^*r*^ were scaled so the inhibitory and excitatory components are equal for *N* = 100, resulting in a collective threshold at that group size that equals the mean individual threshold.

**Figure S7:**
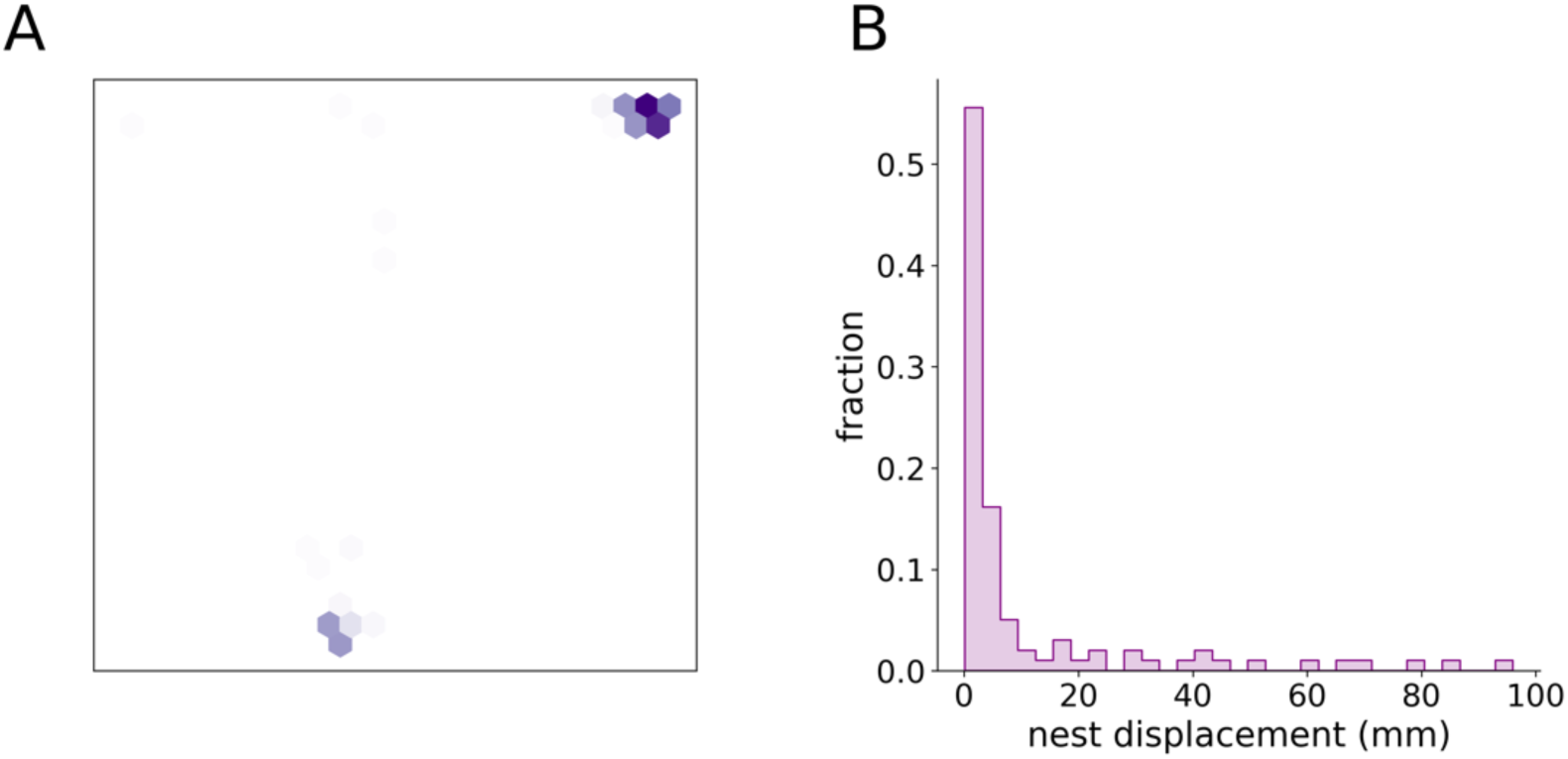
Ants have a tendency to settle at the previous nest site after a perturbation. (**A**) Heatmap depicting the nest locations of one colony of 36 ants throughout a full experiment with variable perturbation amplitudes (>48hrs). (**B**) Histogram of nest displacement distances following full evacuations of the nest in response to temperature perturbations. Data points are pooled across all colonies from the experiment with 36 individually tagged ants per colony and variable perturbation amplitudes. The plot shows a high tendency of colonies to return to their pre-perturbation nest site. This could be explained by a hitherto unknown, weakly volatile pheromone that marks the nest location.

**Video 1: Evacuation response to a strong temperature perturbation**. The video shows a typical nest evacuation of a colony of 36 clonal raider ants in response to a strong (40°C) step increase in temperature. For visualization purposes, the video is background subtracted to show only the behavior of the ants and is sped up x3. The time counter shows the time in seconds relative to the onset of the perturbation.

**Video 2: Aborted response to a mild temperature perturbation**. The video an example for an aborted response of a colony of 36 clonal raider ants in response to a mild (33°C) step increase in temperature. For visualization purposes, the video is background subtracted to show only the behavior of the ants and is sped up x3. The time counter shows the time in seconds relative to the onset of the perturbation.

## References

1. Branco, T. & Redgrave, P. The Neural Basis of Escape Behavior in Vertebrates. Annual Review of Neuroscience 43, 417–439 (2020).

2. Card, G. M. Escape behaviors in insects. Current Opinion in Neurobiology 22, 180–186 (2012).

3. Ter Hofstede, H. M., Schöneich, S., Robillard, T. & Hedwig, B. Evolution of a Communication System by Sensory Exploitation of Startle Behavior. Current Biology 25, 3245–3252 (2015).

4. Davidson, J. D. & El Hady, A. Foraging as an evidence accumulation process. PLOS Computational Biology 15, e1007060 (2019).

5. Levy, S. & Bargmann, C. I. An Adaptive-Threshold Mechanism for Odor Sensation and Animal Navigation. Neuron 105, 534-548.e13 (2020).

6. Fechner, G. T. Elemente der Psychophysik. Breitkopf und Härtel (1860).

7. Swets, J. A. Is there a sensory threshold? Science 134, 168–177 (1961).

8. Green, D. M. & Swets, J. A. Signal detection theory and psychophysics. (John Wiley & Sons, Ltd, 1966).

9. Ori, H., Marder, E. & Marom, S. Cellular function given parametric variation in the hodgkin and huxley model of excitability. Proceedings of the National Academy of Sciences of the United States of America 115, E8211–E8218 (2018).

10. Van Vreeswijk, C. & Sompolinsky, H. Chaos in neuronal networks with balanced excitatory and inhibitory activity. Science 274, 1724–1726 (1996).

11. Van Vugt, B. et al. The threshold for conscious report: Signal loss and response bias in visual and frontal cortex. Science 360, 537–542 (2018).

12. Ferrell, J. E. & Machleder, E. M. The Biochemical Basis of an All-or-None Cell Fate Switch in Xenopus Oocytes. Science 280, 895–898 (1998).

13. Paliwal, S. et al. MAPK-mediated bimodal gene expression and adaptive gradient sensing in yeast. Nature 2006 446:7131 446, 46–51 (2007).

14. Segall, J., Block, S. & Berg, H. Temporal comparisons in bacterial chemotaxis. PNAS 83, 8987–8991 (2009).

15. Biswas, D., Bhattacharya, S. & Iglesias, P. A. Enhanced chemotaxis through spatially-regulated absolute concentration robustness. bioRxiv 2021.07.10.451673 (2021). doi:10.1101/2021.07.10.451673

16. Hölldobler, B. & Wilson, E. The Superorganism: The Beauty, Elegance, and Strangeness of Insect Societies. (W.W. Norton & Co., 2008).

17. Detrain, C. & Deneubourg, J.-L. Self-organized structures in a superorganism: do ants ‘ behave’ like molecules? Physics of Life Reviews 3, 162–187 (2006).

18. Johnson, B. R. & Linksvayer, T. A. Deconstructing the superorganism: Social physiology, groundplans, and sociogenomics. Quarterly Review of Biology 85, 57–79 (2010).

19. Feinerman, O. & Korman, A. Individual versus collective cognition in social insects. Journal of Experimental Biology 220, 73–82 (2017).

20. Bonabeau, E., Theraulaz, G. & Deneubourg, J.-L. Quantitative Study of the Fixed Threshold Model for the Regulation of Division of Labour in Insect Societies. Proceedings of the Royal Society B: Biological Sciences 263, 1565–1569 (1996).

21. Ulrich, Y., Saragosti, J., Tokita, C. K., Tarnita, C. E. & Kronauer, D. J. C. Fitness benefits and emergent division of labour at the onset of group living. Nature 560, 635–638 (2018).

22. Beshers, S. N. & Fewell, J. H. Models of division of labor in social insects. Annual Review of Entomology 46, 413–40 (2001).

23. Robinson, G. E. Regulation of division of labor in insect societies. Annual Review of Entomology 37, 637–665 (1992).

24. Cook, C. N. & Breed, M. D. Social context influences the initiation and threshold of thermoregulatory behaviour in honeybees. Animal Behaviour 86, 323–329 (2013).

25. Ravary, F., Lecoutey, E., Kaminski, G., Châline, N. & Jaisson, P. Individual experience alone can generate lasting division of labor in ants. Current biology : CB 17, 1308–12 (2007).

26. Pratt, S. C. & Sumpter, D. J. T. A tunable algorithm for collective decision-making. Proceedings of the National Academy of Sciences of the United States of America 103, 15906–15910 (2006).

27. Pratt, S. C., Mallon, E. B., Sumpter, D. J. T. & Franks, N. R. Quorum sensing, recruitment, and collective decision-making during colony emigration by the ant Leptothorax albipennis. Behavioral Ecology and Sociobiology 52, 117–127 (2002).

28. Robinson, E. J. H., Franks, N. R., Ellis, S., Okuda, S. & Marshall, J. A. R. A simple threshold rule is sufficient to explain sophisticated collective decision-making. PLoS ONE 6, (2011).

29. Seeley, T. D. et al. Stop signals provide cross inhibition in collective decision-making by honeybee swarms. Science 335, 108–111 (2012).

30. Cole, B. J. Short-Term Activity Cycles in Ants: Generation of Periodicity by Worker Interaction. https://doi.org/10.1086/285156 137, 244–259 (2015).

31. Couzin, I. D. Collective cognition in animal groups. Trends in Cognitive Sciences 13, 36–43 (2009).

32. Bonabeau, E., Theraulaz, G. & Deneubourg, J. L. The synchronization of recruitment-based activities in ants. Biosystems 45, 195–211 (1998).

33. Cole, B. J. & Cheshire, D. Mobile cellular automata models of ant behavior: Movement activity of Leptothorax allardycei. American Naturalist 148, 1–15 (1996).

34. Sasaki, T. & Pratt, S. C. The psychology of superorganisms: collective decision making by insect societies. Annu. Rev. Entomol 63, 259–75 (2018).

35. Reina, A., Bose, T., Trianni, V. & Marshall, J. A. R. Psychophysical Laws and the Superorganism. Scientific Reports 8, 4387 (2018).

36. Chandra, V. et al. Social regulation of insulin signaling and the evolution of eusociality in ants. Science 361, 398–402 (2018).

37. Ulrich, Y. et al. Response thresholds alone cannot explain empirical patterns of division of labor in social insects. PLoS Biology 19, 2020.03.05.963207 (2021).

38. Chandra, V., Gal, A. & Kronauer, D. J. C. Colony expansions underlie the evolution of army ant mass raiding. Proceedings of the National Academy of Sciences 118, (2021).

39. Ofstad, T. A., Zuker, C. S. & Reiser, M. B. Visual place learning in Drosophila melanogaster. Nature 474, 204–207 (2011).

40. Frank, D. D., Jouandet, G. C., Kearney, P. J., MacPherson, L. J. & Gallio, M. Temperature representation in the Drosophila brain. Nature 519, 358–361 (2015).

41. Okada, J. & Toh, Y. Shade Response in the Escape Behavior of the Cockroach, Periplaneta americana. https://doi.org/10.2108/zsj.15.831 15, 831–835 (1998).

42. Gal, A., Saragosti, J. & Kronauer, D. J. C. anTraX, a software package for high-throughputvideo tracking of color-tagged insects. eLife 9, 1–32 (2020).

43. King, A. J. & Cowlishaw, G. When to use social information: The advantage of large group size in individual decision making. Biology Letters 3, 137–139 (2007).

44. Sumpter, D. J. T., Krause, J., James, R., Couzin, I. D. & Ward, A. J. W. Consensus Decision Making by Fish. Current Biology 18, 1773–1777 (2008).

45. Ward, A. J. W., Krause, J. & Sumpter, D. J. T. Quorum decision-making in foraging fish shoals. PLoS ONE 7, (2012).

46. Kao, A. B. & Couzin, I. D. Decision accuracy in complex environments is often maximized by small group sizes. Proceedings of the Royal Society B: Biological Sciences 281, (2014).

47. Torney, C. J., Lorenzi, T., Couzin, I. D. & Levin, S. A. Social information use and the evolution of unresponsiveness in collective systems. Journal of the Royal Society Interface 12, (2015).

48. Mateo, D., Kuan, Y. K. & Bouffanais, R. Effect of Correlations in Swarms on Collective Response. Scientific Reports 7, 1–11 (2017).

49. Laurent Salazar, M. O., Deneubourg, J. L. & Sempo, G. Information cascade ruling the fleeing behaviour of a gregarious insect. Animal Behaviour 85, 1271–1285 (2013).

50. King, A. J. et al. Selfish-herd behaviour of sheep under threat. Current Biology 22, R561–R562 (2012).

51. Granovetter, M. Threshold Models of Collective Behavior. American Journal of Sociology 83, 1420–1443 (1978).

52. Barrat, A., Barthélemy, M. & Vespignani, A. Dynamical Processes on Complex Networks. dpcn (Cambridge University Press, 2008).

53. Gleeson, J. P. Binary-state dynamics on complex networks: Pair approximation and beyond. Physical Review X 3, 021004 (2013).

54. Dorogovtsev, S. N., Goltsev, A. V. & Mendes, J. F. F. Critical phenomena in complex networks. Reviews of Modern Physics 80, 1275 (2008).

55. Jackson, D. E. & Ratnieks, F. L. W. Communication in ants. Current Biology 16, R570–R574 (2006).

56. Razin, N., Eckmann, J.-P. & Feinerman, O. Desert ants achieve reliable recruitment across noisy interactions. Journal of the Royal Society, Interface / the Royal Society 10, 20130079 (2013).

57. Marom, S. Neural timescales or lack thereof. Progress in Neurobiology 90, 16–28 (2010).

58. Wark, B., Lundstrom, B. N. & Fairhall, A. Sensory adaptation. Current Opinion in Neurobiology 17, 423–429 (2007).

59. Hong, S. y Arcas, B., Fairhall, A. L. & Agüera y Arcas, B. Single neuron computation: from dynamical system to feature detector. Neural Computation 19, 3133–3172 (2007).

60. Lundstrom, B. N., Higgs, M. H., Spain, W. J. & Fairhall, A. L. Fractional differentiation by neocortical pyramidal neurons. Nature Neuroscience 11, 1335–1342 (2008).

61. Destexhe, A. & Marder, E. Plasticity in single neuron and circuit computations. Nature 431, 789–795 (2004).

62. Marder, E. Variability, compensation, and modulation in neurons and circuits. Proceedings of the National Academy of Sciences 108, 15542–15548 (2011).

63. Trible, W., McKenzie, S. K. & Kronauer, D. J. C. Globally invasive populations of the clonal raider ant are derived from Bangladesh. Biology Letters 16, 20200105 (2020).

64. Trible, W. et al. orco mutagenesis causes loss of antennal lobe glomeruli and impaired social behavior in ants. Cell 170, 727-735.e10 (2017).

